# Third generation indexing for third generation sequencing

**DOI:** 10.1101/2020.05.07.082347

**Authors:** Abdulqader Jighly

**Affiliations:** Agriculture Victoria, AgriBio, Centre for AgriBiosciences, Bundoora, Victoria 3083, Australia

## Abstract

Indexing of DNA sequences is the art of sorting massive genomic data in a user-friendly structure to enable rapid accessing and comparing of different patterns in the data. Current genome assemblers use general algorithms for string indexing that do not exploit the special structural arrangement of genomes. Here, I am proposing a new algorithm that indexes only the configuration of microsatellite motifs along reads assuming that the order of microsatellites will be the same in overlapped sequences. The index size is >1000 times smaller than currently used indices and it has higher tolerance to the high error rates produced by third generation sequencing platforms. The results showed that the proposed algorithm can rapidly detect overlaps among considerable proportion of uncorrected long reads (~50% of all simulated base pairs with average read size of 8.16 kb and total error rates of 14.4%) to build large initial contigs. Unassembled reads can be then mapped to these contigs or can be assembled with them with currently used algorithms. Thus, the proposed algorithm can efficiently be used as an initial stage to significantly reduce the number of pairwise sequence comparisons among reads and/or references and improve the performance of different software but not replacing them. The algorithm was also useful for comparative genomics and detect large locally colinear blocks and structural variations among ten *saccharomyces cerevisiae* strains. The proposed algorithm has the power to make de novo assembly of individuals as routine activity which can lead to more accurate variant calling and pan genomics.

---

Introduction

Sequence assembly is the reconstruction of the original chromosomal DNA by aligning and merging shorter nucleotide fragments. Assembly can be done with the aid of a reference backbone that is similar, but not identical, to the mapped reads or can be done de novo to reconstruct a new full sequence from scratch. Ideally, full de novo assembly can reveal the actual variation among individuals as mapping reads to a reference genome is biased toward the reference variation, but it is more expensive and current algorithms require lots of resources. Even though the development of reference graphs such as HISAT2 (Kim et al. 2019) allowed for mapping reads to standing variations instead of a single reference genome, which significantly improved the assembly of highly diverged reads, it is impossible to include all possible variants in one graph especially for organisms with inadequate variants database. For example, recent de novo assembling of new human genomes discovered more novel sequences (Chaisson et al. 2015; Seo et al. 2016; Shi et al. 2016; Ameur et al. 2018; Wong et al. 2018) although over 2500 genomes have been sequenced so far (http://www.1000genomes.org).

The first step in assembly is to create an index for the data. Indexing of DNA sequences enabled efficient storing, accessing and processing of high-throughput sequencing data without necessarily traversing all reads. Since their use in genomics, indexing algorithms have rapidly evolved with two major generations. In the first, classical full-text indices such as suffix trees, suffix arrays and directed acyclic word graphs were used (McCreight 1976; Blumer et al. 1985; Manber and Myers 1993), while compressed text indices such as the compressed suffix array and the FM-index (Ferragina and Manzini 2001; Grossi et al. 2003) were used in the second generation to reduce memory and computational burden. Yet, all compression-based indexing methods depend on general string compression algorithms and do not exploit the genomic-special characteristics and structures. For this reason, the size of indices cannot fall below the sequence high-order entropy regardless of the compression method. To speed up mapping of read or even enable de novo assembly as a routine activity, it is important to develop new ultra-compressed indexing algorithms to ease the handling of growing data.

A more contiguous de novo assembly can be achieved now with third-generation sequencing methods (TGS). TGS generate longer sequence reads compared to the first- and second-generation platforms but they produce higher error rate per base. The size of TGS reads is long enough to capture multiple featured sequences such as microsatellites in a single read. Microsatellites are short sequence motifs (1-6 bp) repeated tandemly that were reported in all eukaryotic and most prokaryotic genomes with high and non-random frequency. The order of microsatellites in different overlapping reads is the same. This can be exploited to reduce the complexity of de novo read mapping analysis by indexing the order of microsatellite motifs instead of indexing the whole sequence of the read. This method has the power to compact the complexity of the human genome to a level smaller than the complexity of *E. coli* genome. With enough microsatellite density, it should have more tolerance to the high sequencing error produced by TGS compared to previous indexing methods as I will show later.

## The algorithm

The proposed algorithm is designed to boost current mapping and de novo assemblers but not to replace them. Full read comparison is still required after matching the configuration of microsatellites to ensure to ensure the detection of truly overlapping reads and to find the exact overlapping boundaries among reads. The advantage here is to dramatically reduce the number of all versus all raw read comparisons to few candidate overlaps using an ultra-compressed index for the whole genome/sequencing output. This will facilitate the use of more accurate / time-consuming pairwise sequence alignment algorithms that can accommodate the high error rate of TGS reads given that only few full sequence alignments will be required. The algorithm can be integrated with current software as an initial stage to rapidly build preliminary large contigs with considerable proportion of the reads. The remaining unaligned reads through microsatellite order can be then assembled using traditional algorithms or can be mapped to these initial large contigs. The same indexing algorithm can also be used for whole genome comparison across closely related taxa to detect large syntenic regions having the same configuration of microsatellites. Below, I will provide information on how to best implement such algorithm in practice.

### Binary representation of microsatellites

Representing the four nucleotides without ambiguity can be encoded with 2 bits of data while the 20 amino acids can be represented with 5 bits. In the case of microsatellites, there are 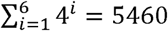 possible permutations for microsatellite motifs ranging from 1 to 6 nucleotides. However, this number can be reduced to 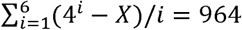 unique motifs. The division on *i* is to account for nucleotide shift (e.g. AAC=ACA=CAA); while *X* represents the overlap between microsatellite motifs of length *i* and shorter motifs due to the within microsatellite tandem repeat (e.g. ACACAC=ACAC=AC). Full equations can be found in text S1. Thus, a single motif can be represented with 10 bits of data if we are indexing microsatellites with a maximum motif length of six nucleotides. Having more bits per unit will not end with a large index size as interval sequences among microsatellites will not be indexed. For example, the human genome has only ~2.7 million microsatellites out of the ~3.2 billion nucleotides.

### Detecting overlapping microsatellite configurations between sequences

Three methods can be used to detect overlaps among sequences using microsatellite configurations:

#### 1. Fixed-size microsatellite *k-mer* matching

Unlike whole sequence indexing, short microsatellite *k-mers* are acceptable due to 1) the large number of possible permutations of microsatellites 964k, where *k* is the *k-mer* size, 2) the small number of microsatellites compared to the number of nucleotides and 3) because matched microsatellite configurations are accompanied with more comprehensive full sequence alignments. However, some microsatellites have higher frequency in the genome compared to others, e.g. monomer (only four types) and hexamer (670 types) microsatellites represent 39% and 2% of all microsatellites in the human genome, respectively. A script to aid the approximation of an appropriate *k-mer* value given the organism’s microsatellite density and bias can be found at text S2. The script counts the most probable *k-mer* events (events with the highest probabilities) for different *k* values, with probabilities sum up to a user specified threshold given an input frequency of each microsatellite motif. As output example, a *4-mer* index would have 964^4^= 8.6 × 10^11^ possible events, but only the most probable 56,018 events will represent 25% of all *4-mers* of the human genome. Thus, for the 2.7 million microsatellites of the human genome, these 56,018 events will theoretically appear 675,000 times with an average of ~12 initial hits on the genome for each one. The number of initial hits can be further reduced after the extension of the hits and getting more matching microsatellites.

This method should be very quick as it only involves direct and continuous matching between read and/or the reference without any amendment. It should have optimal performance with corrected long reads because they can be easily assembled with accurate long *k-mers* and low sequencing depth. When dealing with uncorrected reads, using small *k-mers* is advantageous because the high error rates associated with TGS reads destruct the detection of some microsatellites leading to multiple interrupts of microsatellite orders along the read. Thus, using small *k-mers* will be recommended with caution to avoid ending with huge number of matched microsatellite configurations which may require extensive full pairwise sequence alignments to select the candidate true match. However, the following two methods may be more appropriate in such cases.

#### 2. Probability of the overlaps

As microsatellite motifs vary in their density along the genome, different fixed-size *k-mer* events will have variable probabilities depending on the frequency of each motif in the genome. For example, the sum of all probabilities of all possible hexamer-hexamer *2-mer*s (448,900 events) for the human genome should equal to 4 × 10^−4^ which is much smaller than the sum of all probabilities for all possible *2-mer* mono-motifs (16 events with sum of probabilities = 0.15) or even *8-mer* mono-motifs (65,536 events with sum of probabilities = 5.4 × 10^−4^). Thus, matching microsatellite configurations could be more appropriate when considering the probability of the overlapping microsatellites rather than their counts.

The implementation of this method requires prior knowledge of the occurring probability for different microsatellite motifs in the studied genome or in related organisms. Alternatively, motif probabilities can be derived from their frequencies in all reads assuming that the reads are normally distributed along the genome. The matching stage should start with a *2-mer* or *3-mer* indexing followed by an extension process. The match probability is the multiplication of the occurrence probabilities of all motifs in the overlapped region. Candidate matches are these with probabilities smaller than a user-specified threshold. This process is equivalent to filtering hits based on their e-values with the standard BLAST algorithm.

#### 3. Probability of the overlaps considering the interval distances among microsatellites

The high error rate of TGS reads can alter sequences to distort the detection of existing microsatellites or to form random pseudo microsatellites. Therefore, using a continuous matching of microsatellites on uncorrected reads will end with large proportion of unassembled sequences especially for shorter ones. To solve this issue, the extension of initial hits should ignore some microsatellites on both the subject and the query in a process that mimic the permitting of insertions and deletions in whole sequence alignment. This method requires indexing the distance between each microsatellite and the following one to constrain the search boundaries of the query microsatellite on the subject. Like the previous method, the matching stage should start with a *2-mer* or *3-mer* indexing. During the extension, each adjacent microsatellite (from both sides of the matched *k-mer*) will be compared to all microsatellites on the subject within a specific range. This range differ between mapping reads to a reference and de novo assembly.

When mapping reads to a reference, the first position on the subject to be compared with the query microsatellite should be:

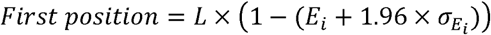

While the last one should be:

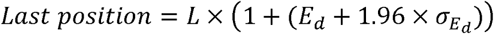

Where *L* is the length of the sequence between the closest <u>*matched*</u> microsatellite and the tested one on the query, *E*_*i*_ is the insertion error rate, *E*_*d*_ is the deletion error rate and *σ* is the standard deviation of error rates. Both *E*_*i*_ and *E*_*d*_ are specific to the applied TGS platform and protocol.

For de novo assembly, both compared reads could have deletions and/or insertions. The range of positions can be given as:

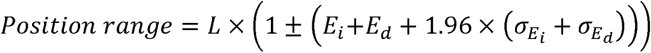

### Testing the algorithm

Two tests were performed. First, simulating long reads and aligning their microsatellites to the original reference sequence. The first 10^5^ base pairs of the Arabidopsis’s chromosome one sequence were used, and 100 long reads was simulated using PaSS software (Zhang et al. 2019). Microsatellites was detected on the simulated reads as well as the original sequence using MISA software with a minimum size of 10 bp and three tandem repeats (Beier et al. 2017). The first and the second detection method was initially used with a *4-mer* and *2-mer* search, respectively, and then the third method was used with a probability threshold of 10^−6^. The second test involved whole genome alignment of ten *Saccharomyces cerevisiae* strains using the configuration of microsatellites to detect locally colinear blocks along their genomes.

#### 1. Mapping of simulated reads

The size of the simulated reads ranged between 0.33 kb and 27.2 kb, covering 94.8% of the whole reference sequence with an average size of 8.16 kb (Figure 1c) and total error rates (insertion, deletion and substitution) of 14.4%. Only 51 simulated reads had more than four microsatellites and they been used for the mapping. Of these, 25 reads with average size of 14.14 kb were truly mapped to the reference (Figure 1a) with the first method covering 83.2% of the reference sequence with an average match length of 4.9 microsatellites (Figure 1d). At low probability threshold, the performance of method two was almost the same as method one without any significant advantage on this specific dataset.

**Figure 1.**
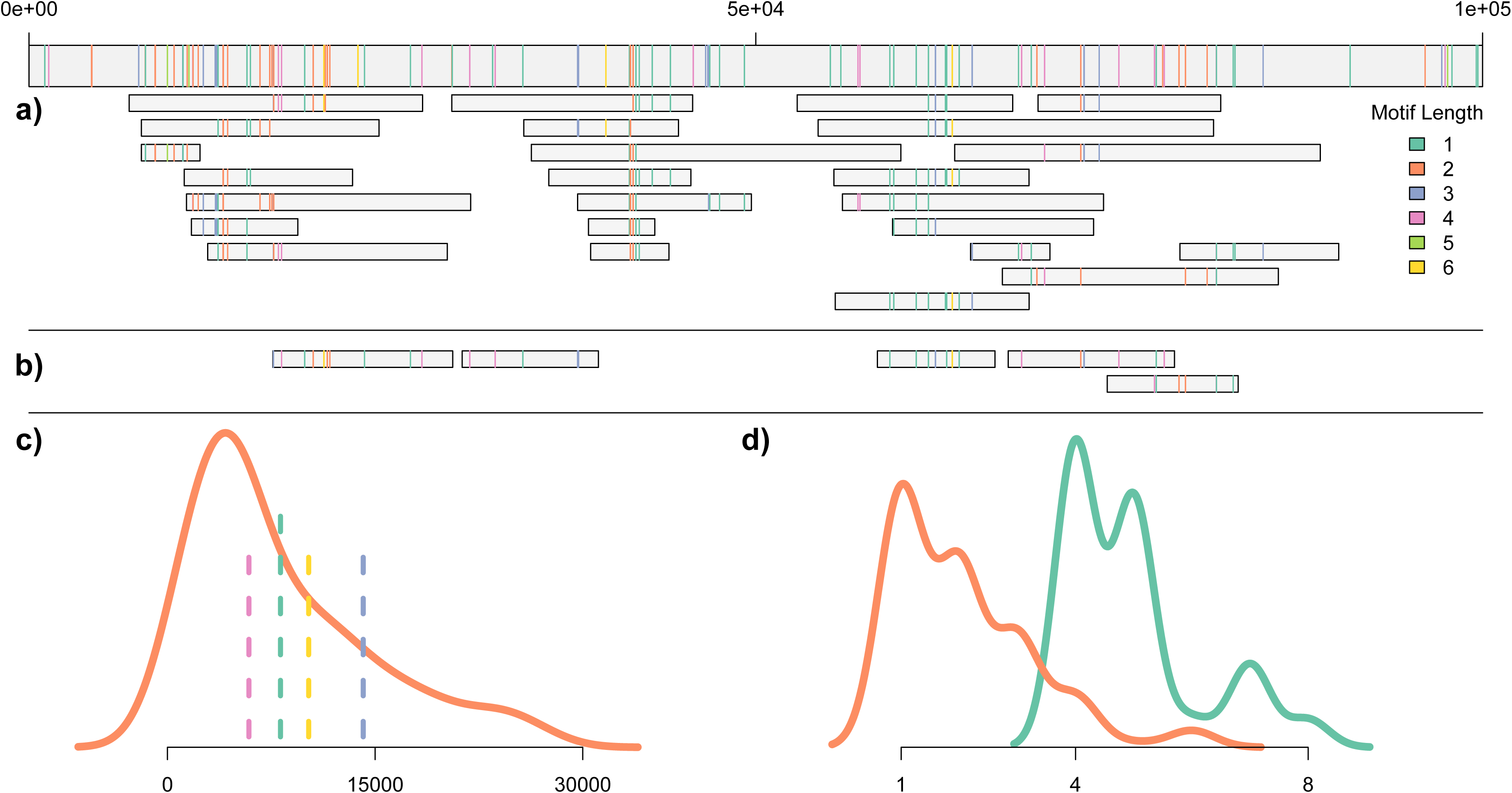
Mapping of simulated reads on the reference sequence. Colored lines on the reference/reads represent the motif size of different microsatellites. a) Mapping uncorrected simulated long reads to the original *Arabidopsis thaliana* sequence using the configurations of their microsatellites; b) extra reads mapped only using method three; c) density plot showing the size distribution of simulated reads, the green vertical line shows the average size of all reads, the blue line shows the average size of reads aligned in (a), the yellow line shows the average size of reads aligned in (b), and the pink line shows the average size of unmapped reads; d) the orange density plot shows the motif size of the matched microsatellites for the mapped reads in (a), while the green density plot shows the number of the matched microsatellite configurations in them.

Five extra reads were further mapped using the third method (Figure 1b) with average size of 10.2 kb while the remaining unmapped reads (70 reads) had an average size of 5.88 kb (Figure 1c). The cumulative size of all mapped reads was 404.5 kb representing 49.6% of the size of all simulated reads and 64% of the reads used in the analysis. As expected, method three performed best as it mapped all reads detected with the other methods and was also more efficient with shorter reads. It performed better in regions with low density of microsatellites because large interval sequences between microsatellites allow for higher chance to create pseudo microsatellites and interrupt the overlapped configurations.

#### 2. Whole genome alignment

A total of 5,440 unique microsatellites were detected across all 10 *saccharomyces cerevisiae* strains of which 63.7% and 9.6% were conserved with the same configurations in all 10 or in 9 strains, respectively, while 9% were strain specific microsatellites. Large structural variations were also detected in which affected sequences had conserved microsatellite configurations regardless of their chromosomal location. For instance, a translocation from the middle of chromosome 17 of the strain P283 to the beginning of chromosome 15 was detected in two fragments with sizes 38.22 kb and 131.3 kb (Figure 2). Another translocation was detected for the same strain with size 69.66 kb from the middle of chromosome 14 to the beginning of chromosome 9. Also, for the line FostersB, a large translocation (145.78 kb) within chromosome 13 was detected while the line Signa1278b had a large deletion (268.28 kb) at the end of chromosome 12 (Table S1). The three methods for detecting overlapped microsatellite configurations performed the same on this dataset due to the high similarity among the 10 strains. However, and like the previous mapping example, it is expected that the third method will be the best for more diverged sequences as more interrupts to microsatellite configurations are expected. The example here proved that locally colinear blocks can be easily detected and the compressed representation of the genome through microsatellites.

**Figure 2.**
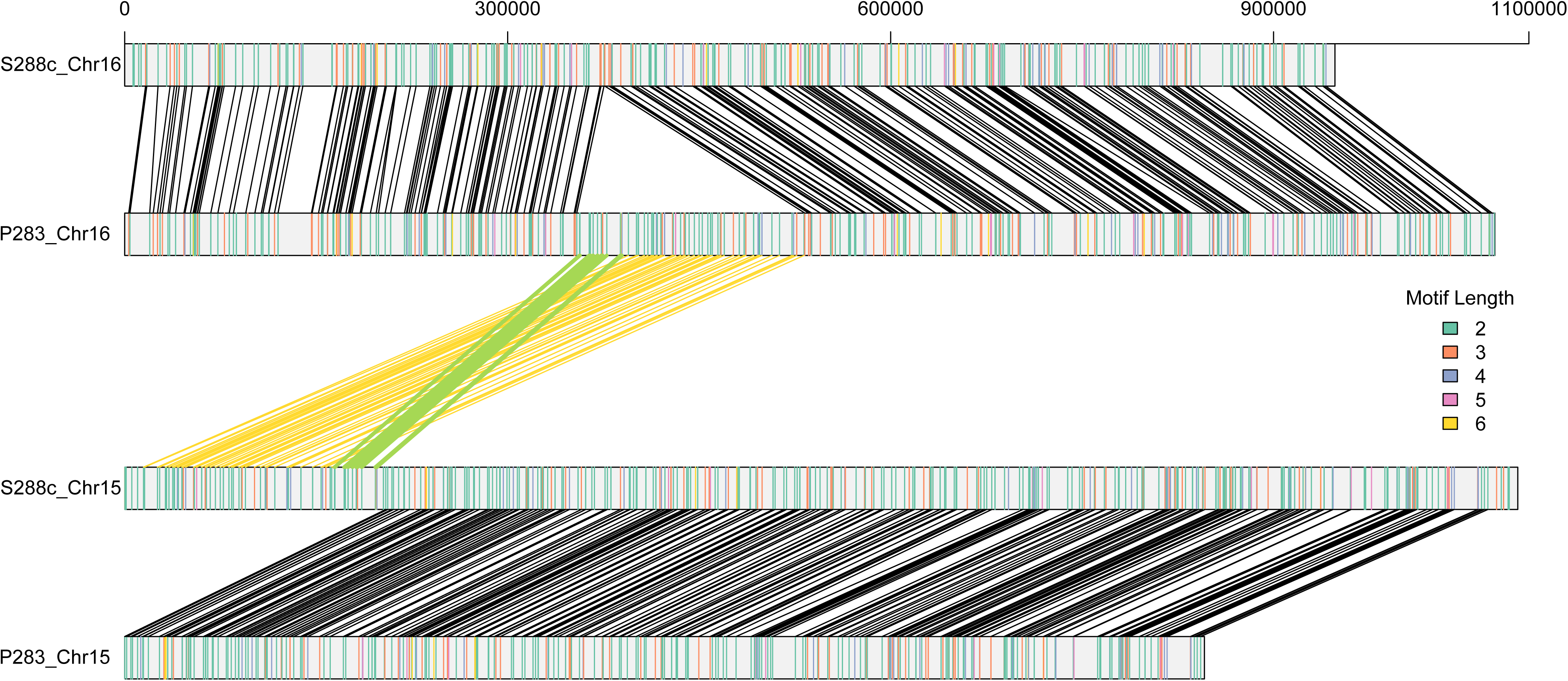
The alignment of microsatellites along the *saccharomyces cerevisiae* chromosomes 15 and 16 for the strains S288c and P283. Colored lines on the chromosomes represent the motif size of different microsatellites. Black lines represent the matched microsatellites configurations. Green and yellow lines represent the translocations from chromosome 16 to 15 in line P283 with sizes 38.22 kb and 131.3 kb, respectively.

### Conclusion

Several algorithms have been proposed to assemble corrected or uncorrected long reads (Chin et al. 2016; Koren et al. 2017; Chin and Khalak 2019; Kolmogorov et al. 2019; Ruan and Li 2020) but still the computational burden and the assembly of uncorrected long reads are major limitations. These tools can be boosted with the algorithm proposed here as an initial step to reduce the complexity of overlapping uncorrelated raw reads. This algorithm detected overlaps among long reads and found large locally colinear blocks by indexing only the order of microsatellites along sequences which is >1,000 times smaller than the whole sequence. The dependency on anchored sequences along the read instead of its whole length can reduce the negative effect of TGS high error rates on sequence assembly. Structural variations can also be detected if they have enough microsatellite density. With such highly efficient complexity reduction algorithm, de novo assembly for thousands of individuals with high sequencing depth can become a routine activity to replace mapping reads to references. Scaffolds of de novo assembled individuals can be compared with the same method to assist with getting a full variant calling and pan genome annotation. Moreover, with some modification, graph-based methods can also be applied on microsatellite configurations to further improve their performance.

## Competing of interested

I declare that I have no competing interests

## Supplementary materials

**Table S1.** The alignment of microsatellite configurations among ten *saccharomyces cerevisiae* strains. Sheet ‘Info’ presents the GeneBank IDs for the used sequences and describes the major detected structural variations. Sheets from one to sixteen representing the 16 *saccharomyces cerevisiae* chromosomes.

**Text S1.** Equations explaining the number of unique microsatellites for each motif size

