## Supplementary material for "Third generation indexing for third generation sequencing": Text S1

Text S1. Equations explaining the number of unique microsatellites for each motif size

| Microsatellite Length (*i*) | Number of Permutations (*p*) | Unique microsatellites (*u*) | Cumulative Unique (*cu*) | Number of Bits | Unique no reverse (*ur*) |
| --- | --- | --- | --- | --- | --- |
| 1 | 4 | 4 | 4 | 2 | 2 |
| 2 | 16 | 6 | 10 | 4 | 4 |
| 3 | 64 | 20 | 30 | 5 | 10 |
| 4 | 256 | 60 | 90 | 7 | 33 |
| 5 | 1024 | 204 | 294 | 9 | 102 |
| 6 | 4096 | 670 | 964 | 10 | 350 |

Column [Number of Permutations (*p*)]:

- *p_i_* = 4*^i^*

Column [Unique microsatellites (*u*)]:

- *u_1_* = 4/1 = 4
- *u_2_* = (*p_2_* – *p_1_*) / 2 = 12/2 = 6
- *u_3_* = (*p_3_* – *p_1_*) / 3 = 60/3 = 20
- *u_4_* = (*p_4_* – *p_2_*) / 4 = 240/4=60
- *u_5_* = (*p_5_* – *p_1_*) / 5 = 1020/5=204
- *u_6_* = (*p_6_* – *p_3_* – *p_2_* + *p_1_*) / 6 = 670

Column [Number of Bits]:

- log_2_(*cu_i_*)

It could also be possible to reduce the number of unique microsatellites by counting each forward microsatellite and its reverse complementary as one unique microsatellite. This step can reduce one bit required for saving each microsatellite and will allow for direct comparison among sequences with both the forward and reverse directions. On the other hand, it increases the probability of each unique microsatellite as it will be the sum of the forward and the reverse complementary motif probabilities. Thus, it might worth explain this from. The number of unique microsatellites assuming forward and reverse complementary directions should be *u* / 2. However, this is applicable only for motifs with odd length. The reverse complementary of some motifs with even length is the same as the forward sequence, e.g. ACGT. Thus, the true number of unique microsatellites considering no reverse complementary can be given with the following equations:

Column [Unique no reverse (*ur*)]:

- *ur_2_* = (*u_2_* + (*u_1_ / 2*)) / 2= 4
- *ur_4_* = (*u_4_* + (*2u_2_ / 2*)) / 2=33
- *ur_6_* = (*u_6_* + (*3u_3_ / 2*)) / 2 = 350
