## Supplementary material for "Third generation indexing for third generation sequencing": Text S2

```
library(gtools)
x=cbind(1:6,c(0.39,0.227,0.075,0.219,0.068,0.02),c(4,6,20,60,204,670))
kmer=2:10
thres=c(seq(0.1,0.99,0.05),0.99)
Events=matrix(NA,nrow=length(kmer),ncol=length(thres))
for (j in 1:length(kmer)) {
	print(j)
	table1=permutations(nrow(x),kmer[j],x[,1],repeats=TRUE,set=F)
	table2=permutations(nrow(x),kmer[j],x[,2],repeats=TRUE,set=F)
	table3=permutations(nrow(x),kmer[j],x[,3],repeats=TRUE,set=F)
	table3=apply(table3,1,prod)#number of possible kmers for each permuation
	table2=apply(table2,1,prod)#product of probs for all permutations
	table4=table2/table3#probability for a single kmer
	table1=cbind(table1,table3,table2,table4)
	table1=table1[order(-table1[,kmer[j]+3]),]
	table1=as.matrix(table1)
	for(i in 2:nrow(table1)) table1[i,kmer[j]+2]=table1[i,kmer[j]+2]+table1[i-1,kmer[j]+2]
	for (i in 1:length(thres)) {
		minID=which(abs((table1[,kmer[j]+2]-thres[i])) == min(abs(table1[,kmer[j]+2]-thres[i])))
		if((table1[minID,kmer[j]+2]-thres[i])>0) {
			Events[j,i]=sum(table1[1:(minID-1),kmer[j]+1])
			f=abs(table1[minID,kmer[j]+2]-thres[i])
			Events[j,i]=Events[j,i]+round(f/table1[minID,kmer[j]+3],0)
		} else {
			Events[j,i]=sum(table1[1:minID,kmer[j]+1])
			f=abs(table1[minID,kmer[j]+2]-thres[i])
			Events[j,i]=Events[j,i]+round(f/table1[minID+1,kmer[j]+3],0)
		}
	}
}
row.names(Events)=kmer
colnames(Events)=thres
write.table(Events,"Events.txt",quote=F,sep='\t')
```
